# LRRC8A regulates hypotonicity-induced NLRP3 inflammasome activation

**DOI:** 10.1101/2020.06.11.146225

**Authors:** Jack P. Green, Tessa Swanton, Lucy V. Morris, Lina Y. El-Sharkawy, James Cook, Shi Yu, James Beswick, Antony Adamson, Neil Humphreys, Richard A. Bryce, Sally Freeman, Catherine Lawrence, David Brough

**Author notes:** Contributed equally (Jack P. Green, Tessa Swanton). To whom correspondence should be addressed: David Brough., Jack Green.

## Abstract

The NLRP3 inflammasome is a multi-molecular protein complex that converts inactive cytokine precursors into active forms of IL-1β and IL-18. The NLRP3 inflammasome is frequently associated with the damaging inflammation of non-communicable disease states and is considered a therapeutic target. However, there is much regarding the mechanism of NLRP3 activation that remains unknown. Chloride efflux is suggested as an important step in NLRP3 activation, but the identity of which chloride channels are involved is still unknown. We used chemical, biochemical, and genetic approaches to establish the importance of Cl^-^ channels in the regulation of NLRP3 activation. Specifically we identify LRRC8A, an essential component of volume-regulated anion channels (VRAC), as a vital regulator of hypotonicity-induced, but not DAMP-induced, NLRP3 inflammasome activation. Although LRRC8A was dispensable for canonical DAMP-dependent NLRP3 activation, this was still sensitive to Cl^-^ channel inhibitors, suggesting there are additional and specific Cl^-^ sensing and regulating mechanisms controlling NLRP3.

## Introduction

Inflammation is an important protective host-response to infection and injury, and yet is also detrimental during non-communicable diseases (1). Inflammasomes are at the heart of inflammatory responses. Inflammasomes are formed by a soluble pattern recognition receptor (PRR), in many cases the adaptor protein ASC (apoptosis-associated speck-like protein containing a CARD), and the protease caspase-1 (2). Inflammasomes form in macrophages in response to a specific stimulus to drive the activation of caspase-1, facilitating the processing of the cytokines pro-interleukin (IL)1β and pro-IL-18 to mature secreted forms, and the cleavage of gasdermin D to cause pyroptotic cell death (2). A number of different inflammasomes have been described, but potentially the inflammasome of greatest interest to non-communicable disease is formed by NLRP3 (NACHT, LRR and PYD domains-containing protein 3) (3). The mechanisms of NLRP3 activation remain poorly understood.

The NLRP3 inflammasome is activated through several routes which have been termed the canonical, non-canonical, and the alternative pathways (3). Activation of the canonical NLRP3 pathway, which has received greatest attention thus far, typically requires two stimuli; an initial priming step with a pathogen associated molecular pattern (PAMP), typically bacterial endotoxin (lipopolysaccharide, LPS), to induce expression of pro-IL-1β and NLRP3, and a second activation step usually involving a damage associated molecular pattern (DAMP), such as adenosine triphosphate (ATP) (4). In 1996, Perregaux and colleagues discovered that hypotonic shock was effective at inducing the release of mature IL-1β when applied to LPS treated human monocytes and suggested the importance of a volume regulated response (5). It was later discovered that hypotonicity induced release of IL-1β via activation of the NLRP3 inflammasome (6), and that this was linked to the regulatory volume decrease (RVD), which is a regulated reduction in cell volume in response to hypo-osmotic-induced cell swelling, and was inhibited by the chloride (Cl^-^) channel blocker NPPB (5-nitro-(3-phenylpropylamino)benzoic acid) (6). The RVD is regulated by the Cl^-^ channel VRAC (volume regulated anion channel). The molecular composition of the VRAC channel was established to consist of an essential LRRC8A sub-unit in combination with other (B-E) LRRC8 sub-units (7, 8). We recently reported that fenamate NSAIDs could inhibit the canonical NLRP3 inflammasome by blocking a Cl^-^ channel, which we suggested could be VRAC (9). We also further characterised the importance of Cl^-^ flux in the regulation of NLRP3, showing that Cl^-^ efflux facilitated NLRP3-dependent ASC oligomerisation (10). Given the poor specificity of many Cl^-^ channel inhibitors, we set out to systematically determine the importance of VRAC and the RVD to NLRP3 inflammasome activation. Using pharmacological and genetic approaches, we discovered that VRAC exclusively regulated RVD dependent NLRP3 activation in response to hypotonicity, and not NLRP3 activation in response to other canonical stimuli. Thus, we provide genetic evidence for the importance of Cl^-^ in regulating NLRP3 via the VRAC dependence of the hypotonicity response, and suggest the presence of additional Cl^-^ sensing mechanisms regulating NLRP3 in response to DAMPs.

## Results

Following publication of the cryo-electron microscopy (cryo-EM) structure of VRAC (11) with inhibitor 4-(2-butyl-6,7-dichloro-2-cyclopentyl-indan-1-on-5-yl)oxobutyric acid (DCPIB) (Figure 1A) (12), we were able to investigate the interaction of established VRAC inhibitors with the channel using molecular modelling. DCPIB was first computationally redocked using Molecular Operating Environment (MOE 2015.08, Chemical Computing Group, Canada) into the homohexameric VRAC structure (PDB code 6NZW, resolution 3.2 Å) (12). The resulting pose produced a reasonable overlay with the cryo-EM conformation of DCPIB, giving a root-mean-square deviation in atomic position of 2.6 Å (Figure 1B, C). DCPIB exhibits an ionic interaction of its carboxylate group with the cationic side-chain of one of the Arg103 residues comprising the electropositive selectivity filter of VRAC (12). Known VRAC inhibitors (9, 13, 14) possessing carboxylic acid groups (flufenamic acid (FFA), mefenamic acid (MFA) and *N-* ((4-methoxy)-2-naphthyl)-5-nitroanthranilic acid (MONNA), the tetrazole moiety (NS3728) and sulfonic acid groups (4-sulfonic calix(6)arene) were then docked into the DCPIB site of VRAC. The most favourably bound poses of these ligands were similarly found to block the pore in a cork-in-bottle manner (12) at the selectivity filter; the ligands’ ionized acidic groups formed strong electrostatic interactions with Arg103 (Figure 1D-H). Tamoxifen, a basic inhibitor of VRAC was also docked into the cryo-EM structure (Figure 1I). Accordingly, tamoxifen docked with its cationic tertiary amino group remote to the Arg103 side-chains (Figures 1l and Supplementary Figure 1); these side-chains instead formed cation-π interactions with the phenyl group of tamoxifen (Figure 1l). The interaction of the tamoxifen pose was computed as having a calculated ligand-binding affinity of -6.5 kcal mol^-1^ via the molecular mechanics/generalized Born volume integration (MM/GBVI) method (15, 16). The binding energies of the anionic ligands were also predicted as favourable, ranging from -5.1 (DCPIB) to -5.8 kcal mol^-1^ (MONNA). This range excludes the larger 4-sulfonic calix(6)arene (calixarene), which gave a binding energy of -9.9 kcal mol^-1^; we note that the GBVI implicit solvent model may be underestimating the high desolvation cost of this polyanionic ligand and therefore overestimating the magnitude of the corresponding binding energy of this compound.

**Figure 1.**
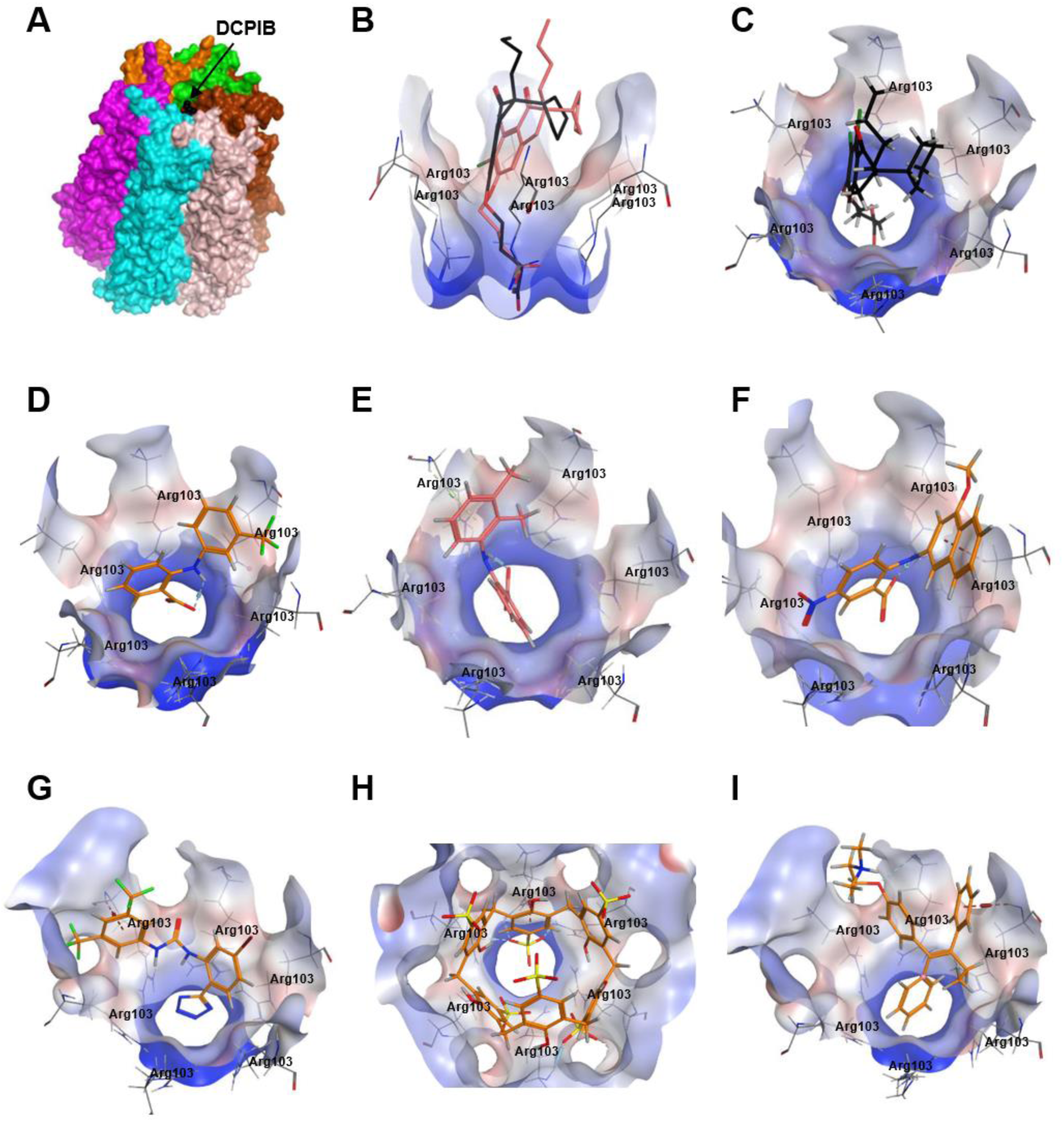
Modelling of proposed VRAC inhibitors on the crystal structure of LRRC8 channels. **(A)** Hexameric cryo-EM structure of VRAC with bound DCPIB (PDB: 6NZW) **(B, D-I)** docked VRAC inhibitors (orange) in the VRAC Arg103 extracellular selectivity filter. Protein surface colored according to hydrophilic (blue), hydrophobic regions (red) and neutral regions (white). **(B)** Side-view of docked DCPIB superimposed on the cryo-EM DCPIB pose (black) **(C)** MM/GBVI binding energies and top-view of cryo-EM pose of DCPIB (black) in VRAC (−5.2 kcal mol^-1^) **(D)** flufenamic acid (−5.1 kcal mol^-1^) **(E)** mefenamic acid (−5.2 kcal mol^-1^) **(F)** MONNA (−5.8 kcal mol^-1^) **(G)** NS3728 (−6.4 kcal mol^-1^) **(H)** 4-sulfonic calix(6)arene (−9.9 kcal mol^-1^) **(I)** tamoxifen (−6.5 kcal mol^-1^).

We then tested the ability of six of the compounds described in Figure 1 (DCPIB, calixarene (Calix), tamoxifen, FFA, MONNA, NS3728) to inhibit hypotonicity-induced VRAC-dependent Cl^-^ flux using the iodide (I^-^) quenching of halide-sensitive YFP H148Q/I152L (17) in live HeLa cells (Figure 2A). In this model, I^-^ enters the cell through open Cl^-^ channels to induce quenching of a mutant EYFP. In response to hypotonic shock to induce VRAC opening, YFP fluorescence was immediately quenched, which was significantly inhibited by tamoxifen (10 µM), MONNA (50 µM), DCPIB (10 µM), FFA (100 µM) and NS3728 (10 µM), but not calixarene (100 µM) (Figure 2A, B). VRAC also regulates RVD in response to cell swelling (7, 8). We measured the RVD by measuring the change in cellular fluorescence in calcein-loaded primary mouse bone marrow-derived macrophages (BMDMs) in response to hypotonicity. Hypotonicity caused a rapid increase in cell volume which declined over time, characteristic of an RVD response (Figure 2C). Similar to the quenching assay, RVD was also significantly inhibited in the presence of tamoxifen, MONNA, DCPIB, FFA and NS3728, but not calixarene (Figure 2C, D). These data suggest that all the molecules in our panel, except calixarene, are bona-fide VRAC inhibitors at the concentrations tested.

**Figure 2.**
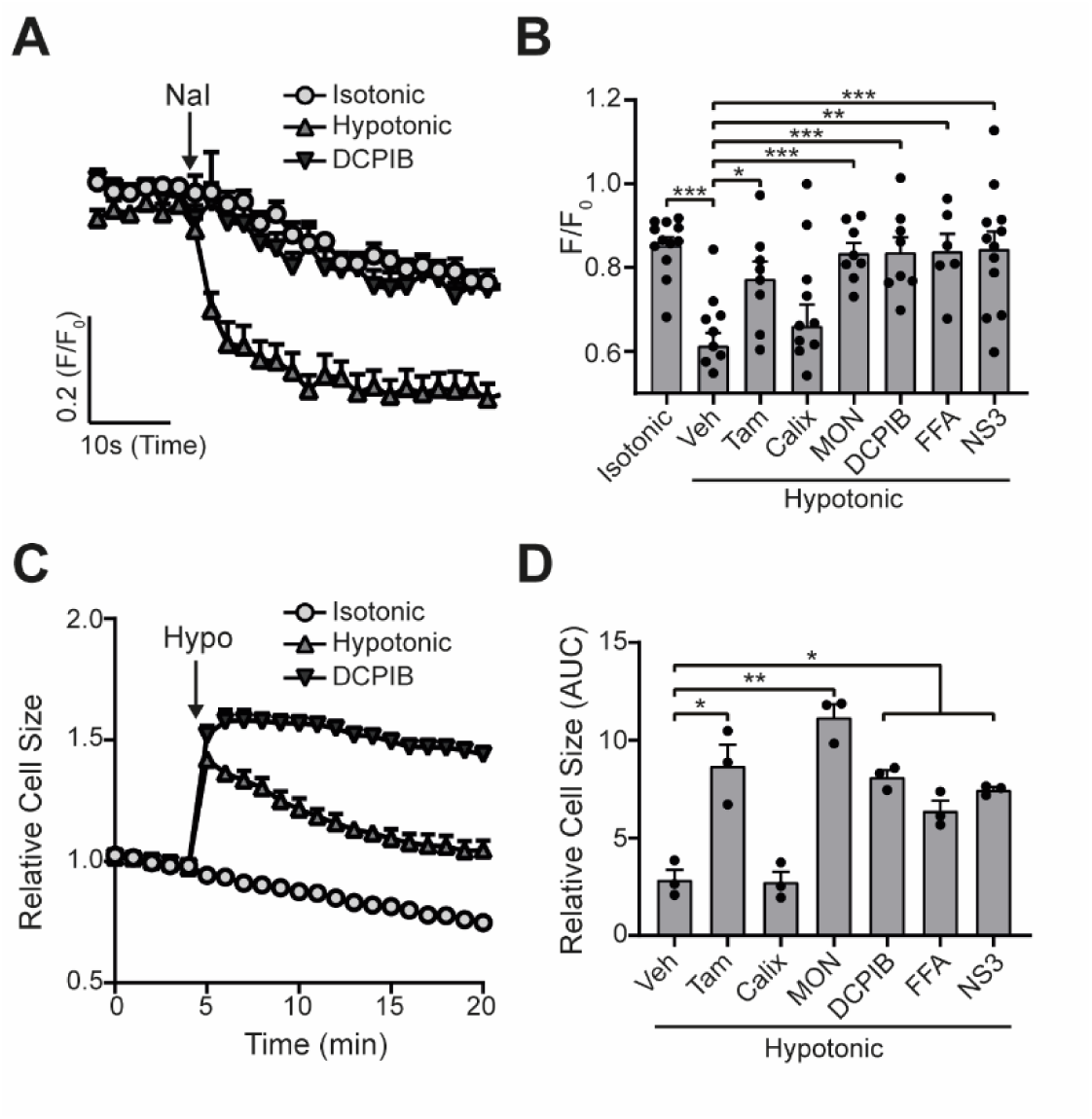
VRAC inhibitors block hypotonicity-induced Cl^-^ channel opening and regulatory volume decrease (RVD). **(A)** Cl^-^ channel opening measured in HeLa cells transiently expressing the halide-sensitive EYFP (H148Q/I152L). HeLa cells were pre-treated with a vehicle control (DMSO) or DCPIB (10 µM) and incubated in an isotonic (310 mOsm kg^-1^) or hypotonic (215 mOsm kg^-1^) solution for 5 minutes before quenching by addition of NaI (40 mM). **(B)** Normalised EYFP (H148Q/I152L) fluorescence values from HeLa cells pre-treated with either a vehicle control (DMSO), tamoxifen (Tam, 10 µM), 4-sulfonic calix(6)arene (Calix, 100 µM), MONNA (MON, 50 µM), DCPIB (10 µM), flufenamic acid (FFA, 100 µM) or NS3728 (NS3, 10 µM). Cells were incubated an isotonic (310 mOsm kg^-1^) or hypotonic (215 mOsm kg^-1^) solution for 5 minutes before quenching by addition of NaI. Fluorescent measurement was taken 30 seconds after NaI addition (n=6-12). **(C)** Relative cell size of murine bone marrow derived macrophages (BMDMs) incubated in isotonic (340 mOsm kg^-1^) or hypotonic (117 mOsm kg^-1^) solution, pre-treated with a vehicle control (DMSO) or DCPIB (10 µM). BMDMs were labelled with the fluorescent dye calcein and area of fluorescence was measured over time. **(D)** Area under the curve of BMDMs incubated in a hypotonic solution (117 mOsm kg^-1^) in the presence of either a vehicle control (DMSO), tamoxifen (Tam, 10 µM), 4-sulfonic calix(6)arene (Calix, 100 µM), MONNA (MON, 20 µM), DCPIB (10 µM), flufenamic acid (FFA, 100 µM) or NS3728 (NS3, 10 µM) (n=3). \**p*<0.05, \*\**p*<0.01, \*\*\**p*<0.01 determined by a one-way ANOVA with Dunnett’s (vs vehicle control) *post hoc* analysis. Values shown are mean plus the SEM.

We tested whether the panel of VRAC inhibitors characterised above could block NLRP3 inflammasome activation and release of IL-1β in response to DAMP stimulation. Primary BMDMs were primed with LPS (1 µg mL^-1^, 4 h), before activation of NLRP3 by ATP (5 mM, 2 h). Inhibitors were given at the same dose they inhibited VRAC above in HeLa cells 15 minutes before the addition of ATP and were then present for the duration of the experiment. Of the panel of verified VRAC inhibitors, only MONNA, FFA and NS3728 consistently inhibited ATP-induced IL-1β release (Figure 3A). At the dose used in this assay, DCPIB did not consistently inhibit ATP-induced IL-1β release (Figure 3A), but at higher concentrations did inhibit NLRP3 activation (Supplementary Figure 2). Likewise, pyroptosis, as measured by LDH release, was significantly reduced by MONNA and FFA, and was unaffected by tamoxifen or calixarene (Figure 3B). The inhibitors that blocked ATP-induced IL-1β release also inhibited ASC oligomerisation, caspase-1 activation, and gasdermin D cleavage (Figure 3C). These data show that some very effective VRAC inhibitors failed to inhibit activation of the NLRP3 inflammasome and release of IL-1β, suggesting that VRAC may not be the molecular target of these molecules inhibiting the inflammasome.

**Figure 3.**
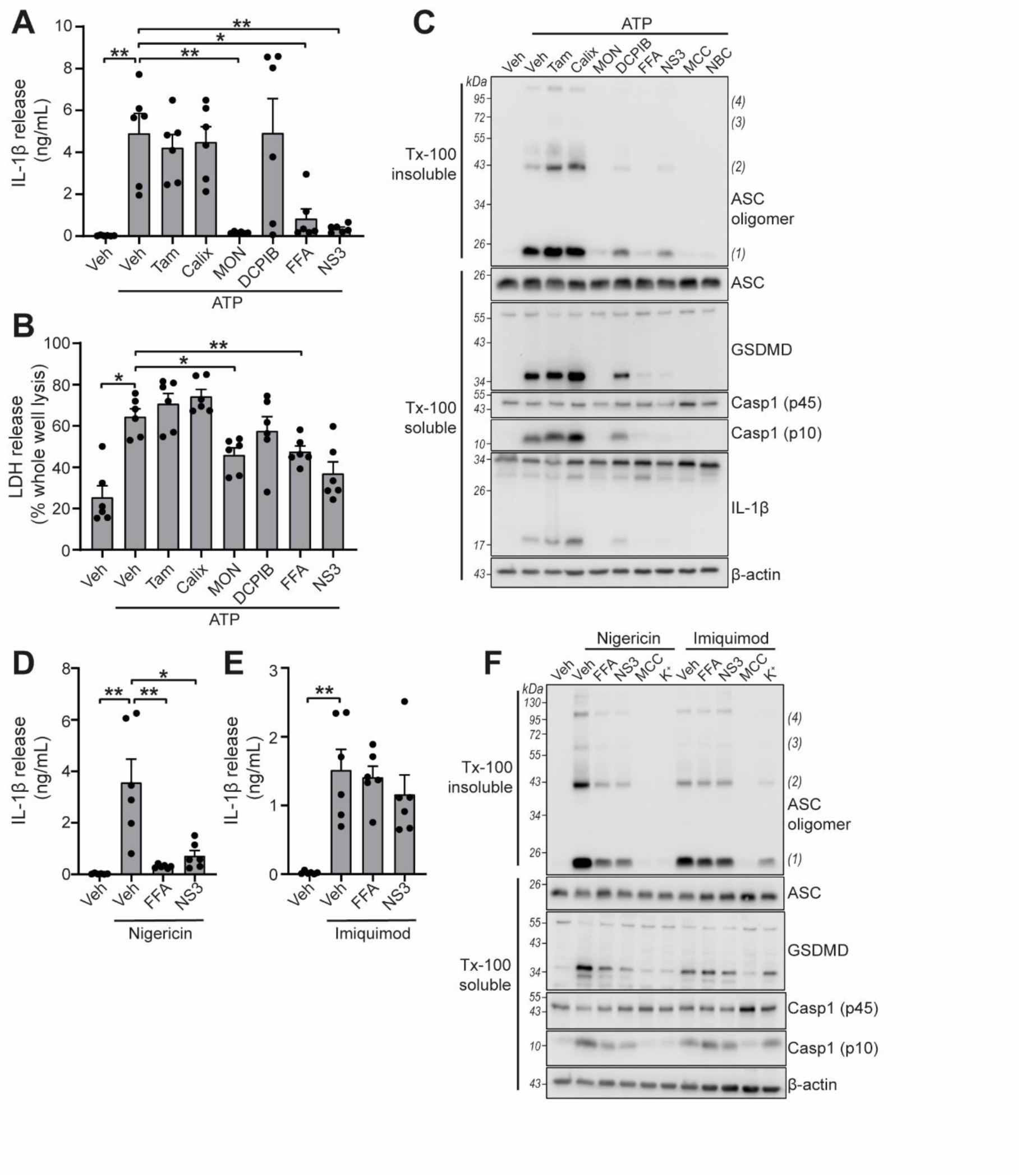
VRAC inhibitors differentially regulate NLRP3. **(A)** IL-1β release was determined by ELISA on supernatants from murine bone marrow derived macrophages (BMDMs). LPS-primed (1 µg mL^-1^, 4 h) BMDMs were pre-treated with either a vehicle control (DMSO), tamoxifen (Tam, 10 µM), 4-sulfonic calix(6)arene (Calix, 100 µM), MONNA (MON, 50 µM), DCPIB (10 µM), flufenamic acid (FFA, 100 µM) or NS3728 (NS3, 10 µM) before stimulation with ATP (5 mM, 2 h) (n=6). **(B)** Cell death determined by an LDH assay of cells treated in (A). **(C)** Western blot of Triton x-100 insoluble crosslinked ASC oligomers and soluble total BMDM cell lysates (cell lysate + supernatant) probed for ASC, GSDMD, caspase-1 and IL-1β. LPS-primed (1 µg mL^-1^, 4 h) BMDMs were pre-treated with either a vehicle control (DMSO), tamoxifen (Tam, 10 µM), 4-sulfonic calix(6)arene (Calix, 100 µM), MONNA (MON, 50 µM), DCPIB (10 µM), flufenamic acid (FFA, 100 µM), NS3728 (NS3, 10 µM) or the NLRP3 inhibitors MCC950 (MCC, 10 µM) and NBC19 (NBC, 10 µM) before stimulation with ATP (5 mM, 2 h) (n=3). **(D)** IL-1β release from LPS-primed (1 µg mL^-1^, 4 h) BMDMs pre-treated with a vehicle control (DMSO), flufenamic acid (FFA, 100 µM) or NS3728 (NS3, 10 µM) before stimulation with nigericin (10 µM, 2 h) (n=6). **(E)** IL-1β release from LPS-primed (1 µg mL^-1^, 4 h) BMDMs pre-treated with a vehicle control (DMSO), flufenamic acid (FFA, 100 µM) or NS3728 (NS3, 10 µM) before stimulation with imiquimod (75 µM, 2 h) (n=6). **(F)** Western blot of Triton x-100 insoluble crosslinked ASC oligomers and soluble total BMDM cell lysates (cell lysate + supernatant) probed for ASC, GSDMD, and caspase-1. LPS-primed (1 µg mL^-1^, 4 h) BMDMs were pre-treated with either a vehicle control (DMSO), flufenamic acid (FFA, 100 µM), NS3728 (NS3, 10 µM), the NLRP3 inhibitor MCC950 (MCC, 10 µM) or KCl (K^+^, 25 mM) before stimulation with nigericin (10 µM, 2 h) or imiquimod (75 µM, 2 h) (n=3). \**p*<0.05, \*\**p*<0.01, determined by a one-way ANOVA with Dunnett’s (vs vehicle control) *post hoc* analysis. Values shown are mean plus the SEM.

ATP-induced NLRP3 activation is dependent upon K^+^ efflux, whereas NLRP3 activation by treatment with the imidazoquinoline compound imiquimod is K^+^ efflux-independent (18). We therefore sought to test if VRAC inhibitors that were effective at blocking ATP-induced inflammasome activation were specific to K^+^ efflux sensitive mechanisms. FFA (100 µM) and NS3728 (10 µM) were effective at blocking IL-1β release after treatment with the K^+^ ionophore nigericin (10 µM, 2 h) (Figure 3D), but were unable to block imiquimod (75 µM, 2 h)-induced IL-1β release (Figure 3E). Similarly, ASC oligomerisation, caspase-1 activation and gasdermin D cleavage induced by nigericin were sensitive to FFA and NS3728 pre-treatment (Figure 3F). However, ASC oligomerisation, caspase-1 activation and gasdermin D cleavage induced by imiquimod were not affected by FFA and NS3728 pre-treatment (Figure 3F). Increased extracellular KCl (25 mM) was sufficient to block nigericin-induced activation, but not imiquimod, demonstrating the K^+^ dependency of nigericin (Figure 3F). These data suggest that these Cl^-^ channel inhibiting compounds exclusively target the K^+^-dependent canonical pathway of NLRP3 activation.

Many Cl^-^ channel inhibiting drugs are known to inhibit multiple Cl^-^ channels, and we established that very effective VRAC inhibitors (tamoxifen and DCPIB) had negligible effect on NLRP3 activation at VRAC inhibiting concentrations. Thus, to conclusively determine the role of VRAC in NLRP3 inflammasome activation we generated a macrophage specific LRRC8A knockout (KO) mouse using CRISPR/Cas9 (Figure 4A). The generation of a macrophage specific LRRC8A KO was required as whole animal LRRC8A KO mice do not survive beyond four weeks and have retarded growth (19). *Lrrc8a*^*fl/fl*^ mice were bred with mice constitutively expressing Cre under the *Cx3cr1* promoter, as previously shown to be expressed in monocyte and macrophage populations (20). This generated mice with the genotype *Lrrc8a*^*fl/fl*^:*Cx3cr1*^*cre*^ (KO) with littermates *Lrrc8a*^*fl/fl*^:*Cx3cr1*^*WT*^ (WT). Cell lysates were prepared from BMDMs and peritoneal macrophages isolated from WT and KO mice and were western blotted for LRRC8A confirming that *Lrrc8a* KO cells were knocked out for LRRC8A (Figure 4B). Functional loss of LRRC8A was confirmed using the calcein RVD assay described above. BMDMs were subjected to a hypotonic shock and changes in calcein fluorescence measured over time. In WT cells there was a characteristic RVD (Figure 4C). However, in *Lrrc8a* KO cells there was complete loss of the RVD response (Figure 4C, D). The absence of RVD was also strikingly evident by observation of the cells by phase contrast microscopy (Figure 4E, Supplementary videos 1, 2). These data confirm functional KO of the VRAC channel in macrophages.

**Figure 4.**
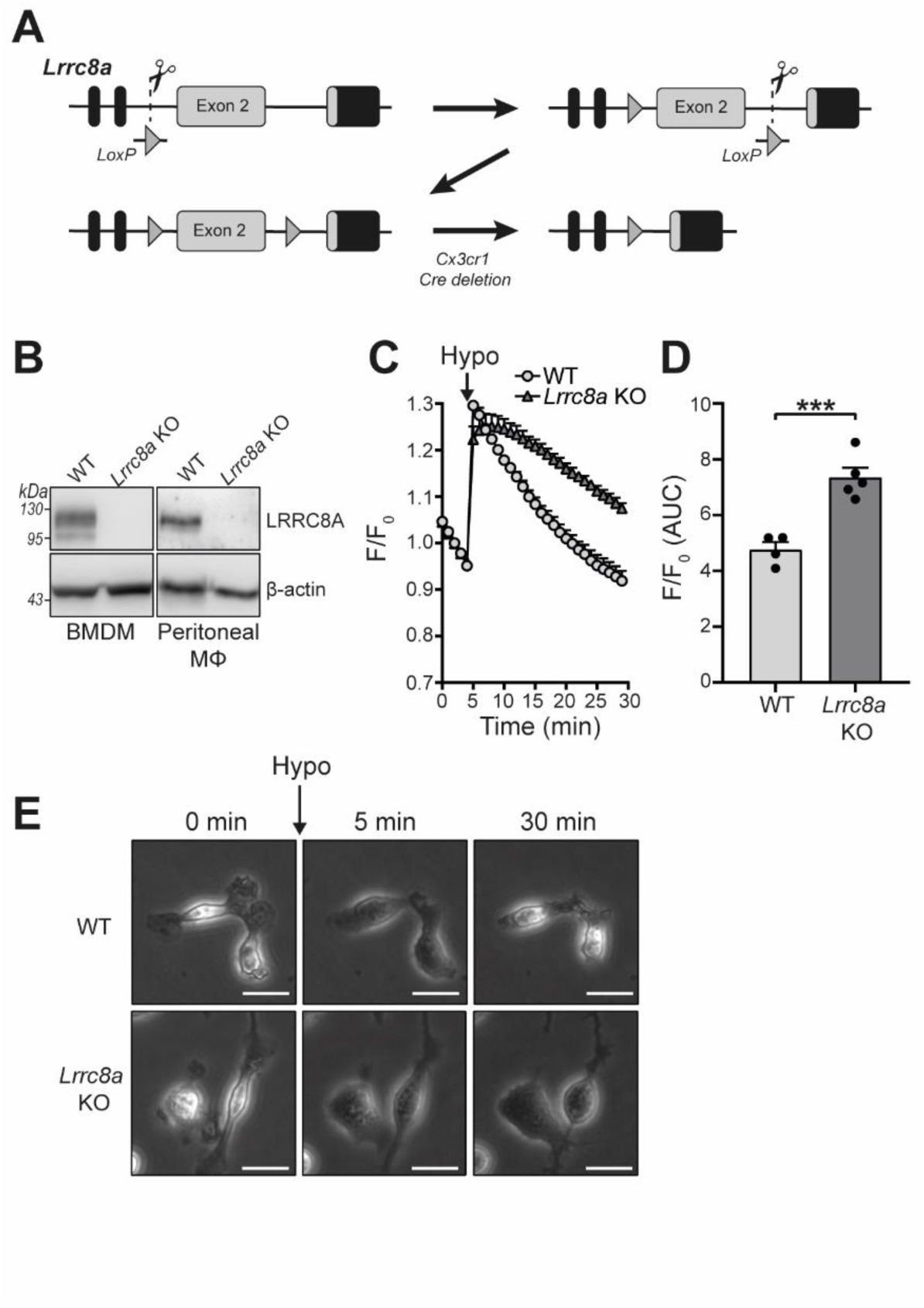
*Lrrc8a* KO macrophages are unable to undergo hypotonicity-induced regulatory volume decrease (RVD). **(A)** Generation of LRRC8A conditional allele. LRRC8A is found on mouse chromosome 2 and consists of 4 exons. Untranslated sequences are represented by black boxes, and coding sequences by grey boxes. Exon 2 contains the vast majority of coding sequence and was flanked by loxP sites in two sequential steps, first integrating the 5’ LoxP by CRISPR-Cas9 (scissors) mediated double strand break and the supply of a homology flanked ssODN repair template containing the loxP site (grey triangle). This 5’ fl background was then bred to establish a colony and the process repeated to integrate the second 3’ loxP on this background. At each step integration of loxP was confirmed by PCR and Sanger sequencing. Finally crossing with a Cre driver knocked into the *Cx3cr1* locus results in recombinase mediated excision of Exon 2. **(B)** Western blot of LRRC8A from wild-type (WT) or *Lrrc8a* knockout (KO) bone marrow derived macrophages (BMDMs) and peritoneal macrophages (Mϕ) (n=3). **(C)** Regulatory volume decrease measured by calcein fluorescence in WT or *Lrrc8a* KO BMDMs incubated in a hypotonic buffer (117 mOsm kg^-1^) (n=4-5). **(D)** Area under the curve (AUC) analysis of (C) (n=4-5). **(E)** Representative phase contrast images of WT or *Lrrc8a* KO BMDMs incubated in a hypotonic buffer (117 mOsm kg^-1^) at indicated time points (n=3, Scale = 20 µm). \*\*\**p*<0.001 determined by an unpaired *t*-test. Values shown are mean plus the SEM.

We next used the *Lrrc8a* KO macrophages to test the hypothesis that VRAC and the RVD were important for NLRP3 inflammasome activation and IL-1β release in response to DAMP stimulation. WT BMDMs and *Lrrc8a* KO BMDMs were primed with LPS (1 µg mL^-1^, 4 h) and then treated with the NLRP3 inflammasome activators ATP (5 mM, 2 h), nigericin (10 µM, 2 h), silica (300 µg mL^-1^, 2 h), or imiquimod (75 µM, 2 h). Knocking out LRRC8A had no effect on the release of IL-1β (Figure 5A) or cell death (Figure 5B). We then used western blotting to determine ASC oligomerisation and caspase-1 activation. In response to the NLRP3 inflammasome activators nigericin, ATP, and imiquimod, there was no effect of LRRC8A KO on ASC oligomerisation or caspase-1 activation (Figure 5C). Furthermore, IL-1β release in response to ATP or nigericin was still inhibited by the VRAC inhibitors flufenamic acid (FFA, 100 µM), and NS3728 (10 µM) in the *Lrrc8a* KO BMDMs, confirming that these inhibitors are inhibiting NLRP3 inflammasome activation by a VRAC-independent mechanism (Figure 5D). Flufenamic acid and NS3728 also inhibited ASC oligomerisation and caspase-1 activation as determined by western blot in the *Lrrc8a* KO BMDMs to the same extent as in the WT (Figure 5E). We then used a murine peritonitis model described previously (9) to investigate the role of LRRC8A *in vivo*. First, we tested if the VRAC inhibitor NS3728 was effective at blocking NLRP3 *in vivo*. Wild type C57BL6/J mice were injected intraperitoneally with NS3728 (50 mg kg^-1^), the NLRP3 inhibitor MCC950 (50 mg kg^-1^), or vehicle control, at the same time as LPS (1 µg, 4 h). NLRP3 was then further activated by intraperitoneal injection of ATP (100 mM, 500 µL, 15 min) and IL-1β release was measured by ELISA of the peritoneal lavage (Figure 5F) and plasma (Figure 5G). Addition of ATP induced a significant increase in the release of IL-1β into peritoneal lavage and this was inhibited by MCC950 and NS3728, indicating an NLRP3-dependent response. IL-6 levels were unaltered in both peritoneal lavage and plasma by addition of NS3728 (Supplementary Figure 3A, B). These data show that NS3728 was able to inhibit NLRP3 *in vivo*. To determine the role of VRAC in this model, we repeated this experiment in our macrophage *Lrrc8a* KO mice and their littermate controls. Macrophage *Lrrc8a* KO mice exhibited normal proportions of myeloid cells in the peritoneum as assessed by flow cytometry (Supplementary Figure 3C-G). Loss of macrophage LRRC8A had no effect on the IL-1β levels in the peritoneal lavage in response to LPS and ATP (Figure 5H), or in the plasma (Figure 5I). Moreover, similar to our *in vitro* findings, NS3728 was still effective at inhibiting this response in the absence of LRRC8A (Figure 5H, I). These data suggested that VRAC was dispensable for NLRP3 activation by DAMP stimulation, and that the VRAC inhibitors are effective at inhibiting NLRP3 in the absence of VRAC, suggesting the presence of another target.

**Figure 5.**
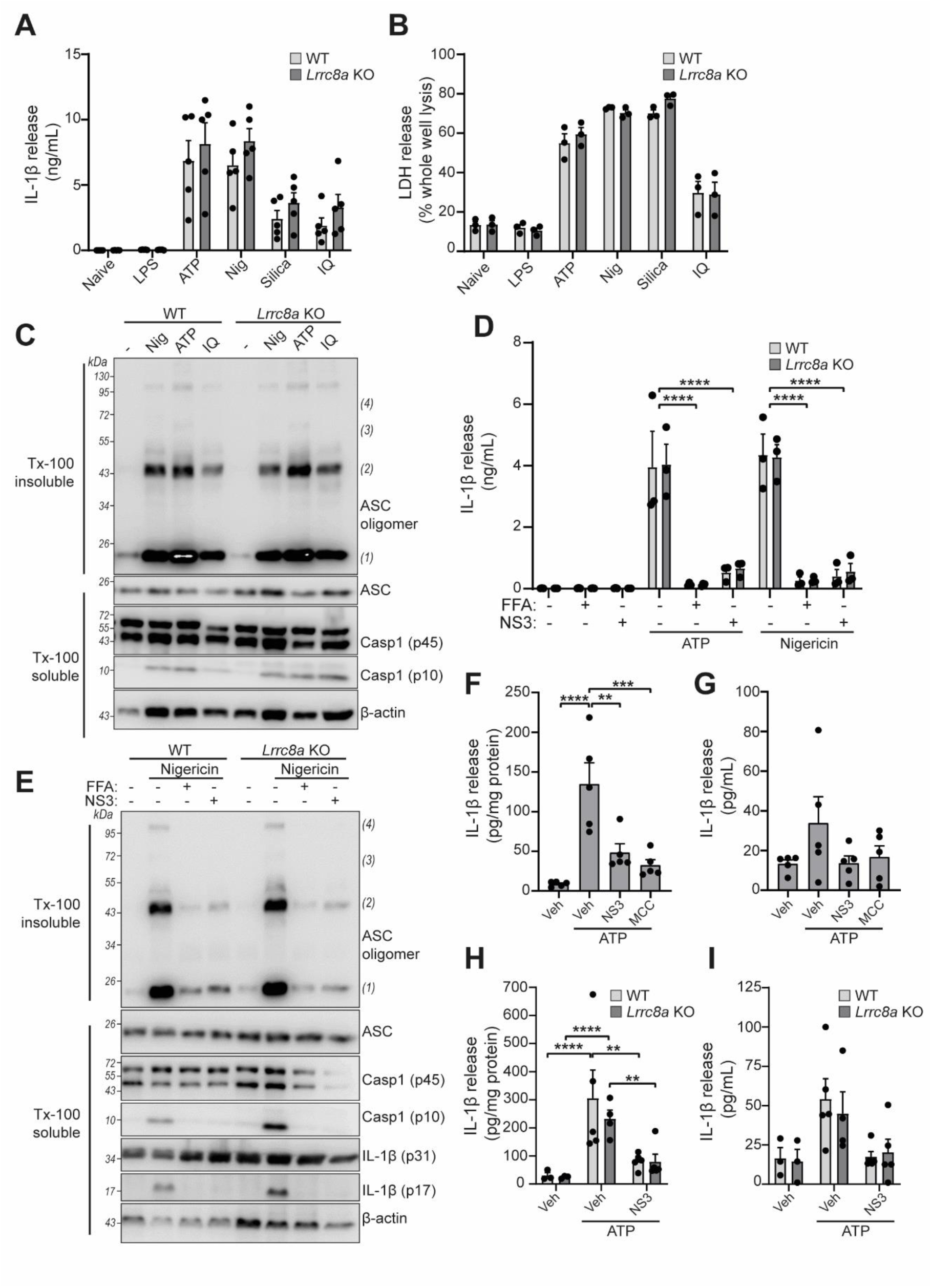
LRRC8A is dispensable for activation of the NLRP3 inflammasome. **(A)** IL-1β release was determined by ELISA on supernatants from wild-type (WT) or *Lrrc8a* knockout (KO) bone marrow derived macrophages (BMDMs). Naïve or LPS-primed (1 µg mL^-1^, 4 h) BMDMs were stimulated with either vehicle, ATP (5 mM), nigericin (Nig, 10 µM), silica (300 µg mL^-1^) or imiquimod (IQ, 75 µM) for 2 h (n=5). **(B)** Cell death determined by an LDH assay of cells treated in (A) (n=3). **(C)** Western blot of Triton x-100 insoluble crosslinked ASC oligomers and soluble total BMDM cell lysates (cell lysate + supernatant) probed for ASC and caspase-1. LPS-primed (1 µg mL^-1^, 4 h) WT or *Lrrc8a* KO BMDMs were stimulated with either nigericin (Nig, 10 µM), ATP (5 mM) or imiquimod (IQ, 75 µM) for 2 h (n=3). **(D)** IL-1β release from LPS-primed (1 µg mL^-1^, 4 h) WT or *Lrrc8a* KO BMDMs pre-treated with a vehicle control (DMSO), flufenamic acid (FFA, 100 µM) or NS3728 (NS3, 10 µM) and then stimulated with ATP (5 mM) or nigericin (10 µM) for 2 h (n=3). **(E)** Western blot of Triton x-100 insoluble crosslinked ASC oligomers and soluble total BMDM cell lysates (cell lysate + supernatant) probed for ASC, caspase-1 and IL-1β. LPS-primed (1 µg mL^-1^, 4 h) WT or *Lrrc8a* KO BMDMs were pre-treated with a vehicle control, flufenamic acid (FFA, 100 µM) or NS3728 (NS3, 10 µM) and stimulated with nigericin (10 µM, 2 h) (n=5). **(F-G)** IL-1β detected by ELISA in the peritoneal lavage (F) or plasma (G) from WT mice. Mice were pre-treated intraperitoneally (i.p.) with a vehicle control, NS3728 (NS3, 50 mg kg^-1^) or MCC950 (MCC, 50 mg kg^-1^) and LPS (1 µg). 4 h after injection with LPS, mice were anaesthetised and injected with additional vehicle control, NS3728 (NS3, 50 mg kg^-1^) or MCC950 (MCC, 50 mg kg^-1^) before i.p. injection of ATP (100 mM, 500 µL, 15 min) (n=5). **(H-I)** IL-1β detected by ELISA in the peritoneal lavage (H) or plasma (I) from *Lrrc8a* KO and WT littermates as treated in (F) (n=3-5). \*\**p*<0.01, \*\*\**p*<0.001, \*\*\*\**p*<0.0001 determined by a one-way ANOVA with Dunnett’s (vs vehicle control) *post hoc* analysis (F,G) or a two-way ANOVA with Tukey’s *post hoc* analysis (A,B,D,H,I). Values shown are mean plus the SEM.

RVD in response to hypo-osmotic-induced cell swelling is documented as an inducer of NLRP3 inflammasome activation and IL-1β release (5, 6). Since *Lrrc8a* KO BMDMs could no longer control their volume in response to hypotonic shock, we tested whether NLRP3 inflammasome activation by hypotonicity was altered. LPS-primed (1 µg mL^-1^, 4 h) BMDMs were incubated in a hypotonic solution (4 h) which caused IL-1β and LDH release from WT cells, and which was significantly inhibited in *Lrrc8a* KO BMDMs (Figure 6A,B). There was no difference in IL-1β release or cell death between ATP-stimulated WT and KO BMDMs (Figure 6A, B). Caspase-1 cleavage and IL-1β processing induced by hypotonicity were also completely inhibited in the absence of LRRC8A (Figure 6C), indicating the response was completely dependent on both NLRP3 and VRAC. Moreover, hypotonicity-induced ASC oligomerisation was also dependent on VRAC (Figure 6D). These data show that in response to hypo-osmotic stress, VRAC was essential for NLRP3 inflammasome activation.

**Figure 6.**
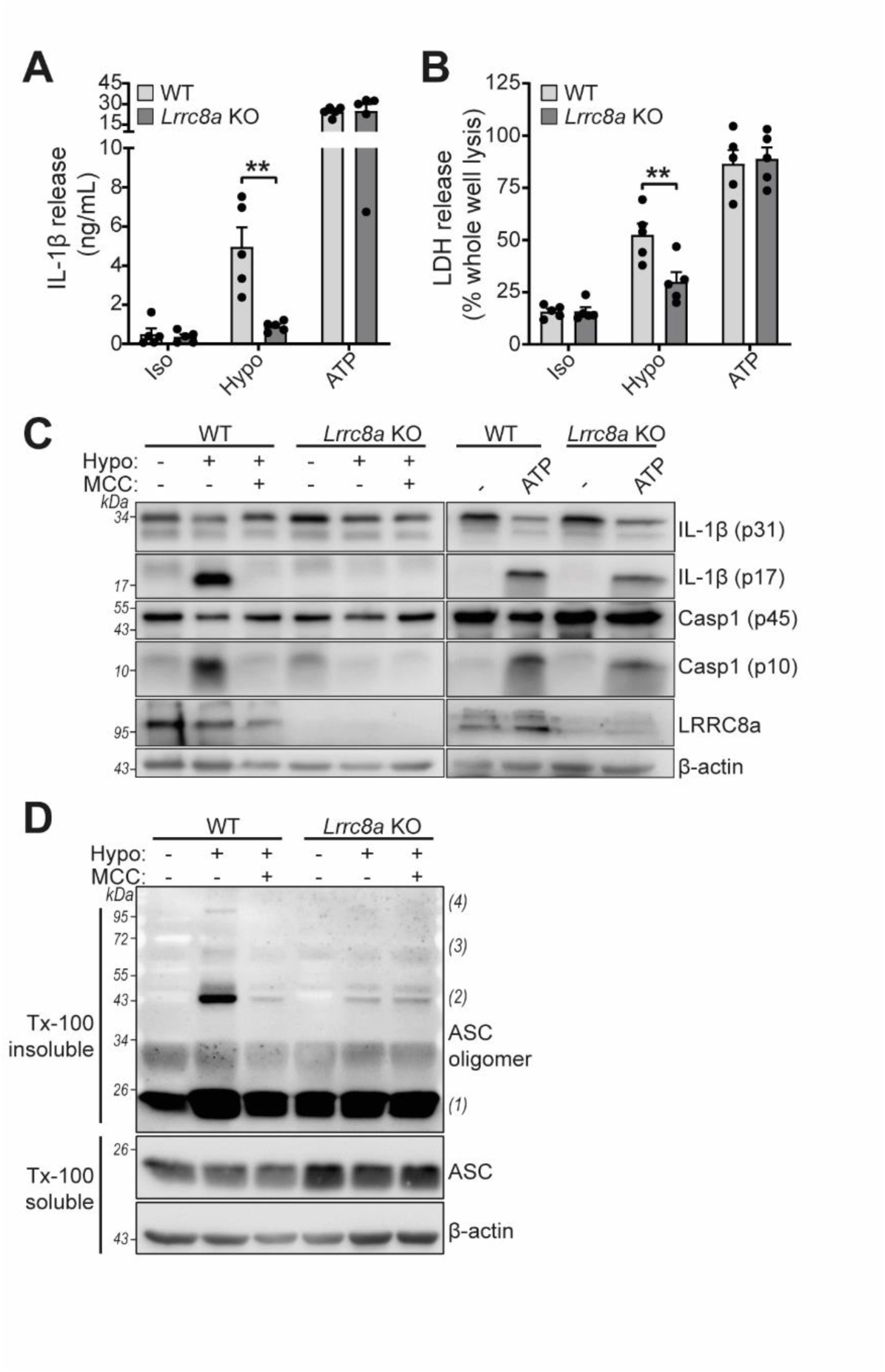
LRRC8a is an essential component of hypotonicity-induced NLRP3 activation. **(A)** IL-1β release was determined by ELISA on supernatants from wild-type (WT) or *Lrrc8a* knockout (KO) bone marrow derived macrophages (BMDMs). LPS-primed (1 µg mL^-1^, 4 h) wild type (WT) or *Lrrc8a* knockout (KO) BMDMs were incubated in either an isotonic buffer (340 mOsm kg^-1^), hypotonic buffer (117 mOsm kg^-1^) or isotonic buffer with ATP (5 mM) for 4 h (n=5). **(B)** Cell death determined by an LDH assay of cells treated in (A). **(C)** Western blot of total BMDM cell lysates (cell lysate + supernatant) probed for IL-1β, caspase-1 and LRRC8a. LPS-primed (1 µg mL^-1^, 4 h) WT or *Lrrc8a* KO BMDMs were pre-treated with a vehicle control or a NLRP3 inhibitor MCC950 (MCC, 10 µM) before incubation in either an isotonic buffer (340 mOsm kg^-1^), hypotonic buffer (117 mOsm kg^-1^) or isotonic buffer with ATP (5 mM) for 4 h. **(D)** Western blot of Triton x-100 insoluble crosslinked ASC oligomers and soluble total BMDM cell lysates (cell lysate + supernatant) probed for ASC. LPS-primed (1 µg mL^-1^, 4 h) WT or *Lrrc8a* KO BMDMs were pre-treated with a vehicle control or MCC950 (MCC, 10 µM) and incubated in either an isotonic buffer (340 mOsm kg^-1^), hypotonic buffer (117 mOsm kg^-1^) for 4 h (n=3). \*\**p*<0.01 determined by a two-way ANOVA with Tukey’s *post hoc* analysis. Values shown are mean plus the SEM.

## Discussion

Pharmacological and biochemical evidence supporting an important role of Cl^-^ ions in the activation of the NLRP3 inflammasome has been provided by various studies over the years (5, 6, 9, 10, 21), although conclusive genetic evidence has been lacking. The promiscuous nature of many Cl^-^ channel inhibiting drugs, and an unresolved molecular identity of major Cl^-^ channels, have prevented the emergence of conclusive genetic proof. However, the discovery that the Cl^-^ channel regulating the RVD (VRAC) was composed of LRRC8 sub-units, and that LRRC8A was essential for channel activity, offered us the opportunity to investigate the direct importance of VRAC in the regulation of NLRP3. Hypotonicity induces cell swelling which is corrected by the VRAC-dependent RVD (7, 8). The RVD was previously linked to NLRP3 activation (6). Thus by knocking out LRRC8A, and thus VRAC, we would discover that VRAC was essential for RVD-induced NLRP3 inflammasome activation, providing strong evidence for the direct requirement of a Cl^-^ channel in NLRP3 inflammasome activation.

However VRAC was only essential for RVD induced NLRP3 activation and was not involved in the NLRP3 response to DAMP stimulation. The fact that our VRAC channel inhibiting drugs block DAMP-induced NLRP3 activation suggests that additional Cl^-^ channels are involved in coordinating NLRP3 responses to other stimuli. Chloride intracellular channel proteins (CLICs 1-6) form anion channels and regulate a variety of cellular processes (22, 23). Localisation of CLIC1 and 4 to membrane fractions in macrophages is increased by LPS stimulation, and RNAi knockdown of both CLIC1 and 4 impaired LPS and ATP-induced IL-1β release from macrophages (24). In addition to CLICs 1 and 4, CLIC5 is also implicated in NLRP3-dependent IL-1β release (25). Knockdown of CLICs 1, 4, and 5 inhibits NLRP3 inflammasome activation in response to the soluble agonists ATP and nigericin, and also the particulate DAMP monosodium urate crystals (25). Thus, it appears that multiple Cl^-^ channels encode diverse signals arising from DAMP stimulation, or from altered cellular homeostasis, to trigger NLRP3 inflammasome activation.

Inhibiting the NLRP3 inflammasome has become an area of intense research interest due to the multiple indications of its role in disease (3). The inhibitor MCC950 is now thought to bind directly to NLRP3 to cause inhibition (26, 27), although it has also been reported to inhibit Cl^-^ flux from macrophages treated with nigericin (28), and was found to bind directly to CLIC1 (29), so it is possible that some of its inhibitory activity may be attributable to an effect on Cl^-^. We found that Cl^-^ channel inhibition blocked IL-1β release in a NLRP3-dependent model of peritonitis, and previously reported protective effects of the fenamate NSAIDs in rodent models of Alzheimer’s disease that we attributed to an effect on Cl^-^ channel inhibition (9). Thus, it is possible that targeting Cl^-^ channels offers an additional route to inhibit NLRP3-dependent inflammation in disease.

In summary, we have reported that hypotonicity induced NLRP3 inflammasome activation depends exclusively on the Cl^-^ channel VRAC, and that different Cl^-^ sensing and regulating systems coordinate the activation of NLRP3 in response to DAMPs. This opens the possibility of discrete Cl^-^ regulating mechanisms conferring selectivity and information about the nature of the NLRP3 activating stimulus. Thus, this investigation has opened the door to further studies on Cl^-^ regulation of NLRP3 and identified the possibility of selective therapeutic intervention strategies informed by the nature of the disease or DAMP stimulus, potentially minimising complications of immunosuppression caused by a blanket NLRP3 inhibition.

## Methods

### Computational chemistry

Docking of ligands to VRAC employed the recently solved cryo-EM structure of VRAC (PDB: 6NZW, resolution 3.2 Å) (12). Tautomeric and ionization states of VRAC amino acid residues at pH 7.4 were assigned using MOE (MOE 2015.08, Chemical Computing Group, Canada). Similarly, ligands were modelled in their ionized forms according to physiological conditions. Docking was performed with the Triangle Matcher placement method of MOE using the London dG scoring function (MOE 2015.08, Chemical Computing Group, Canada). The pocket into which the VRAC inhibitors were docked was that occupied by the (S)-isomer of DCPIB in the cryo-EM structure of VRAC. Rescoring of poses used the molecular mechanics (MM)/generalized Born/volume integral (GBVI) potential (15).

### Antibodies and Reagents

Specific antibodies were used targeting: IL-1β (AF-401-NA, R&D), caspase-1 (ab179515, abcam), gasdermin D (ab20945, abcam), ASC (67824, CST), LRRC8A (8H9, Santa Cruz) and β-actin (A3854, Sigma).

Lipopolysaccharide (LPS, *in vitro: Escherichia coli 026:B6*, L2654, *in vivo: Escherichia coli 0127:B8*, L3880), adenosine triphosphate (ATP, A2383), nigericin (N7143), tamoxifen (T5648), flufenamic acid (FFA, F9005), CP-456773 (MCC950, PZ0280), sodium iodide (NaI, 409286) were purchased from Sigma. Disuccinimidyl suberate (DSS, 21555), calcein AM (C1430) and 4-sulfonic calix[6]arene (10494735) were purchased from Thermo Fisher. 4-[(2-Butyl-6,7-dichloro-2-cyclopentyl-2,3-dihydro-1-oxo-1H-inden-5-yl)oxy]butanoic acid (DCPIB, 1540) was from Tocris. NS3728 and NBC19 were made at the University of Manchester.

### Cell Culture

Primary bone marrow derived macrophages (BMDMs) and peritoneal macrophages were isolated from male and female wild-type C57BL6/J mice. Bone marrow was harvested from both femurs, red blood cells were lysed and resulting marrow cells were cultured in 70% DMEM (10% v/v FBS, 100 U/mL penicillin, 100 μg/mL streptomycin) supplemented with 30% L929 mouse fibroblast-conditioned media for 6-7 days. BMDMs were seeded out the day before at a density of 1×10^6^ mL^-1^ in DMEM (10% v/v FBS, 100 U mL^-1^ penicillin, 100 μg mL^-1^ streptomycin). Peritoneal macrophages were isolated by peritoneal lavage and seeded out overnight at a density of 1×10^6^ mL^-1^ in DMEM (10% v/v FBS, 100 U mL^-1^ penicillin, 100 μg mL^-1^ streptomycin). HeLa cells were seeded out at 0.1×10^6^ mL^-1^ in DMEM (10% v/v FBS, 100 U mL^-1^ penicillin, 100 μg mL^-1^ streptomycin).

### Inflammasome activation assays

Primary BMDMs were primed with LPS (1 µg mL^-1^, 4 h) in DMEM (10% v/v FBS, 100 U mL^-1^ penicillin, 100 μg mL^-1^ streptomycin). After priming, the media was replaced with serum-free DMEM, or when specified an isotonic buffer (132 mM NaCl, 2.6 mM KCl, 1.4 mM KH_2_PO_4_, 0.5 mM MgCl_2_, 0.9 mM CaCl_2_, 20 mM HEPES, 5 mM NaHCO_3_, 5 mM Glucose, pH 7.3, 340 mOsm/kg) or hypotonic buffer (27 mM NaCl, 0.54 mM KCl, 0.3 mM KH_2_PO_4_, 0.5 mM MgCl_2_, 0.9 mM CaCl_2_, 20 mM HEPES, 5 mM NaHCO_3_, 5 mM Glucose, pH 7.3, 117 mOsm kg^-1^). When used, VRAC inhibitors were added 15 minutes before stimulation of the NLRP3 inflammasome.

For analysis of IL-1β release and pyroptosis, cell supernatants were collected. IL-1β release was determined by ELISA (DuoSet, R&D Systems) according to the manufacturer’s instructions. Cell death was assessed by lactate dehydrogenase (LDH) release using CytoTox 96 nonradioactive cytotoxicity assay (Promega) according to manufacturer’s instructions. For western blotting, total cell lysates were made by directly adding protease inhibitor cocktail and Triton x-100 (1% v/v) to each well containing cells and supernatant.

### ASC oligomerisation assay

1 × 10^6^ primary BMDMs were seeded out overnight into 12-well plates. After LPS priming (1 µg mL^-1^, 4 h), cells were incubated in either serum-free DMEM, an isotonic or a hypotonic buffer (as described above) and stimulated as described. BMDMs were lysed directly in-well by addition of protease inhibitor cocktail and Triton x-100 (1% v/v) and lysed on ice. Total cell lysates were then spun at 6800*xg* for 20 min at 4°C to separate the lysate into Triton x-100 soluble and insoluble fractions. The Triton x-100 insoluble fraction (pellet) was then chemically crosslinked by addition of disuccinimidyl suberate (DSS, 2 mM, 30 min, RT) in PBS. Following crosslinking, the insoluble fraction was spun at 6800*xg* for 20 min and the resulting pellet was resuspended and boiled in 40 µL 1X Laemlli buffer. The Triton x-100 soluble fraction was concentrated by trichloroacetic acid (TCA) precipitation. Triton x-100 soluble lysate was mixed 1:1 with TCA (20% w/v) and spun at 14000*xg* for 10 min at 4°C. The pellet was then washed in acetone, spun at 14000*xg* for 10 min at 4°C, and resuspended in 2X Laemlli buffer.

### Regulatory volume decrease (RVD) assay

5 × 10^4^ BMDMs were seeded out into black walled 96-well plates overnight. Cells were loaded with calcein (10 µM, 1 h, 37°C) in an isotonic buffer (132 mM NaCl, 2.6 mM KCl, 1.4 mM KH_2_PO_4_, 0.5 mM MgCl_2_, 0.9 mM CaCl_2_, 20 mM HEPES, 5 mM NaHCO_3_, 5 mM Glucose, pH 7.3, 340 mOsm kg^-1^). Following loading, BMDMs were washed three times with isotonic buffer before incubation with VRAC inhibitors or vehicle control at indicated concentrations for 5 min. GFP fluorescence was then imaged for a further 5 min before hypotonic shock was induced by a five-fold dilution with a hypotonic buffer (0.9 mM CaCl_2_, 20 mM HEPES, 5 mM NaHCO_3_, 5 mM Glucose, pH 7.3), resulting in a final osmolarity of 117 mOsm kg^-1^. GFP fluorescence was measured on an Eclipse Ti inverted microscope (Nikon) and analysed using Image J software. Point visiting was used to allow multiple positions to be imaged within the same time-course and cells were maintained at 37°C and 5% CO_2_. For experiments with VRAC inhibitors, combined treatment with hypotonicity and VRAC inhibitors resulted in some cells undergoing lytic cell death over the course of the experiment and loss of calcein fluorescence. Therefore, GFP fluorescence was used to identify the area of living cells.

### Iodide YFP quenching assay

HeLa cells were seeded at a density of 0.1×10^6^ ml^-1^ in black-walled, clear bottom 96-well plates (Greiner). Transient transfection with the halide-sensitive YFP mutant pcDNA3.1 EYFP H148Q/I152L, a gift from Peter Haggie (Addgene plasmid # 25872), was performed using Lipofectamine 3000 (Thermo Fisher). 18-24 h post-transfection, HeLa cells were washed twice with isotonic buffer (140 mM NaCl, 5 mM KCl, 20 mM HEPES, pH 7.4, 310 mOsm kg^-1^) before 5 min incubation in 50 µL isotonic buffer containing either drug at indicated concentrations, or vehicle. 50 µl isotonic or hypotonic (5 mM KCl, 20 mM HEPES, 90 mM mannitol, pH 7.4, 120 mOsm kg^-1^) buffer containing either drug or vehicle was then added and cells were incubated for a further 5 min. NaI (200 mM, 25 µL) was then added directly to the well, and fluorescence readings were take every 2 seconds using the FlexStation3 plate reader.

### Generation of *Lrrc8a*^*fl/fl*^ mice

We used CRISPR-Cas9 to generate the floxed LRRC8A allele on C57BL/6J background. LRRC8A is a 4 exon gene spanning 26kb on mouse chromosome 2. Only 2 of these exons contain coding sequence, with exon 3 harbouring > 85% of the coding sequence and possessing large introns, and thus an ideal candidate for floxing. We initially attempted the 2-sgRNA, 2-oligo approach described previously (30), but failed to obtain mice with both loxP integrated on the same allele (31). Instead, a colony from a single founder with the 5’ LoxP integrated was established, bred to homozygosity, and used as a background to integrate the second 3’ loxP. For both steps, we used the Sanger WTSI website (http://www.sanger.ac.uk/htgt/wge/, (32)) to design sgRNA that adhered to our criteria for off target predictions (guides with mismatch (MM) of 0, 1 or 2 for elsewhere in the genome were discounted, and MM3 were tolerated if predicted off targets were not exonic). sgRNA sequences were purchased as crRNA oligos, which were annealed with tracrRNA (both oligos supplied IDT; Coralville, USA) in sterile, RNase free injection buffer (TrisHCl 1mM, pH 7.5, EDTA 0.1mM) by combining 2.5 µg crRNA with 5 µg tracrRNA and heating to 95°C, which was allowed to slowly cool to room temperature.

For 5’ targeting the sgRNA GTCTAGTTAGGGACTCCTGG-*GGG* was used, with the ssODN repair template 5’-tccttgacttgctgtttaccgctctcttccccacaccacagttatccacaggaagttacccataacctccctcgtgcacccctacccccaATAACTTCGTATAGCATACATTATACGAAGTTATGGTACCggagtccctaactagacctgctgtctctccatagccctgtctacacct-3’, where capitals indicate the LoxP sequence with a KpnI site, and lower case the homology arms. For embryo microinjection the annealed sgRNA was complexed with Cas9 protein (New England Biolabs) at room temperature for 10 min, before addition of ssODN (IDT) donor (final concentrations; sgRNA 20 ng µL^-1^, Cas9 protein 20 ng µL^-1^, ssODN 50 ng µL^-1^). CRISPR reagents were directly microinjected into C57BL6/J (Envigo) zygote pronuclei using standard protocols (33). Zygotes were cultured overnight and the resulting 2 cell embryos surgically implanted into the oviduct of day 0.5 post-coitum pseudopregnant mice. Potential founder mice were identified by extraction of genomic DNA from ear clips, followed by PCR using primers that flank the homology arms and sgRNA sites (Geno F1 tcagatggcgaaccagaagtc and Geno R1 tacaatgtagtcaggtgtgacg). WT sequences produced a 833bp band, and loxP knock in 873bp, which is also susceptible to KpnI digest. Pups with a larger band were reserved, the band isolated and amplified using high fidelity Phusion polymerase (NEB), gel extracted and subcloned into pCRblunt (Invitrogen). Colonies were mini-prepped and Sanger sequenced with M13 Forward and Reverse primers, and aligned to predicted knock-in sequence. Positive pups were bred with a WT C57BL6/J to confirm germline transmission and a colony established. To integrate the 3’ LoxP we used sgRNA ACTACCCCATTACCTCTTGG-*TGG* with the ssODN repair template 5’-gagggccaaaactgtggaaagcaacacccttgaagtgtaggtggcccctgtgcaccagctctgtgtgtgactgcaaagcccccaccaagaATAACTTCGTATAGCATACATTATACGAAGTTATGGTACCggtaatggggtagttagacgggctgagggcagagcacttgtgtggctt-3’. Again, capitals indicate the LoxP sequence with a KpnI site, and lower case the homology arms.

For this second round of targeting we generated embryos from the LRRC8A-5’fl colony by IVF using homozygous LRRC8A-5’fl mice, and used electroporation (Nepa21 electroporator, Sonidel) to deliver the sgRNA:Cas9 RNP complex and ssODN to the embryos, AltR crRNA:tracrRNA:Cas9 complex (200 ng μL^-1^; 200 ng μL^-1^; 200 ng μL^-1^ respectively) and ssDNA HDR template (500 ng μL^-1^) (34). Zygotes were cultured overnight and the resulting 2 cell embryos surgically implanted into the oviduct of day 0.5 post-coitum pseudopregnant mice. Potential founder mice were identified by extraction of genomic DNA from ear clips, followed by PCR using primers that flank the homology arms and sgRNA sites (Geno F2 atccccactgcttttctgga and Geno R2 ccactcaagagccagcaatg). WT sequences produced a 371bp band, and loxP knock in 411bp, which is also susceptible to KpnI digest. As before, Pups with a larger band were reserved, the band isolated and amplified using high fidelity Phusion polymerase (NEB), gel extracted and subcloned into pCRblunt (Invitrogen). Colonies were mini-prepped and Sanger sequenced with M13 Forward and Reverse primers, and aligned to predicted knock-in sequence. Positive pups were bred with a WT C57BL6/J to confirm germline transmission and a colony established (*Lrrc8a*^*fl/fl*^*). Lrrc8a*^*fl/fl*^ mice were bred with *Cx3cr1*^*cre*^ mice (as previously described (20)) to generate *Lrrc8a*^*fl/fl*^ *Cx3cr1*^*cre/*+^ mice, which specifically induce removal of LRRC8A in cells expressing CX3CR1 (*Lrrc8a KO)*. CX3CR1-cre mice were obtained from a breeding colony at the University of Manchester managed by John Grainger. Experiments using *Lrrc8a* KO cells were compared to wild-type littermate controls (*Lrrc8a*^*fl/fl*^ *Cx3cr1*^+*/*+^).

### In vivo peritoneal inflammation model

All procedures were performed with appropriate personal and project licenses in place, in accordance with the Home Office (Animals) Scientific Procedures Act (1986), and approved by the Home Office and the local Animal Ethical Review Group, University of Manchester. 8-10 week-old male wild type (WT) C57BL/6J mice (Charles River) were used to test efficacy of NS3728 and MCC950. Mice were treated intraperitoneally with LPS (2 µg mL^-1^, 500 µL) and either a vehicle control (5% DMSO (v/v), 5% Cremophor (v/v), 5% ethanol (v/v) in PBS), NS3728 (50 mg kg^-1^) or MCC950 (50 mg kg^-1^) for 4 h. Mice were then anesthetised with isofluorane (induced at 3% in 33% O_2_, 67% NO_2_, maintained at 1–2%) before injection with a vehicle control, NS3728 or MCC950 as before and ATP (0.5 mL, 100 mM in PBS, pH 7.4) or PBS for 15 min. The peritoneum was then lavaged with RPMI 1640 (3 mL) and blood was collected via cardiac puncture. For experiments using *Lrrc8a* knockout (KO) mice, 8-10 week-old male and female *Lrrc8a* KO and WT littermate controls were used and treated as described above. Plasma and lavage fluid was used for cytokine analysis. BCA analysis was performed on the peritoneal lavage fluid to normalise cytokine release to total protein level. Murine studies were performed with the researcher blinded to genotype and treatment for the duration of the experiment.

### Flow cytometry

8-10 week-old male and female *Lrrc8a* KO and WT littermate controls were anesthetised with isofluorane (induced at 3% in 33% O_2_, 67% NO_2_, maintained at 1–2%) and the peritoneal cavity was lavaged with 6 mL RPMI (3% v/v FBS, 100 U mL^-1^ penicillin, 100 μg mL^-1^ streptomycin, 1 mM EDTA) before sacrifice. Red blood cells were lysed using Pharmlyse (BD Biosciences) in H_2_O. Cells were surface stained with fluorescence-conjugated anti-CD45, anti-CD11b, anti-Ly6G, anti-MHCII, anti-F4/80, anti-Ly6C and anti-CX3CR1 antibody cocktail containing Fc block (anti-CD16/CD32) and Tris-EDTA (1 mM). Cells were then fixed (10 min, room temperature) with paraformaldehyde (2% w/v). Live/Dead Fixable Blue stain was used to exclude dead cells. Samples were analysed on an LSRII flow cytometer (Becton-Dickinson) and cell populations characterised as follows:-neutrophils (CD45^hi^/CD11b^hi^/Ly6G^hi^), monocyte-derived-macrophages (CD45^hi^/CD11b^hi^/Ly6G^-^/MHCII^hi^/F4/80^-^), resident macrophages (CD45^hi^/CD11b^hi^/Ly6G^-^/F4/80^hi^), Ly6C^hi^ monocytes (CD45^hi^/CD11b^hi^/Ly6G^-^ /MHCII^-^/F4/80^-^/CX3CR1^hi^/Ly6C^hi^), using FlowJo software.

### Quantification and statistical analysis

Data are presented as mean values plus the SEM. Accepted levels of significance were *P < 0.05, **P < 0.01, ***P < 0.001, ****P < 0.0001. Statistical analyses were carried out using GraphPad Prism (version 8). Data where comparisons were made against a vehicle control, a one way ANOVA was performed with a Dunnett’s *post hoc* comparison was used. Experiments with two independent variables were analysed using a two-way ANOVA followed by a Tukey’s *post hoc* corrected analysis. Equal variance and normality were assessed with Levene’s test and the Shapiro–Wilk test, respectively, and appropriate transformations were applied when necessary. n represents experiments performed on individual animals or different passages for experiments involving HeLa cells.

## Acknowledgments

This work was supported by Medical Research Council Grants MR/N029992/1 and MR/T0116515/1 (to D.B.), The Alzheimer’s Society AS-PhD-16-002 (to D.B.) and by a Presidential Fellowship (University of Manchester, to J.G.).

## Figure Legends

**Supplementary Figure 1.**
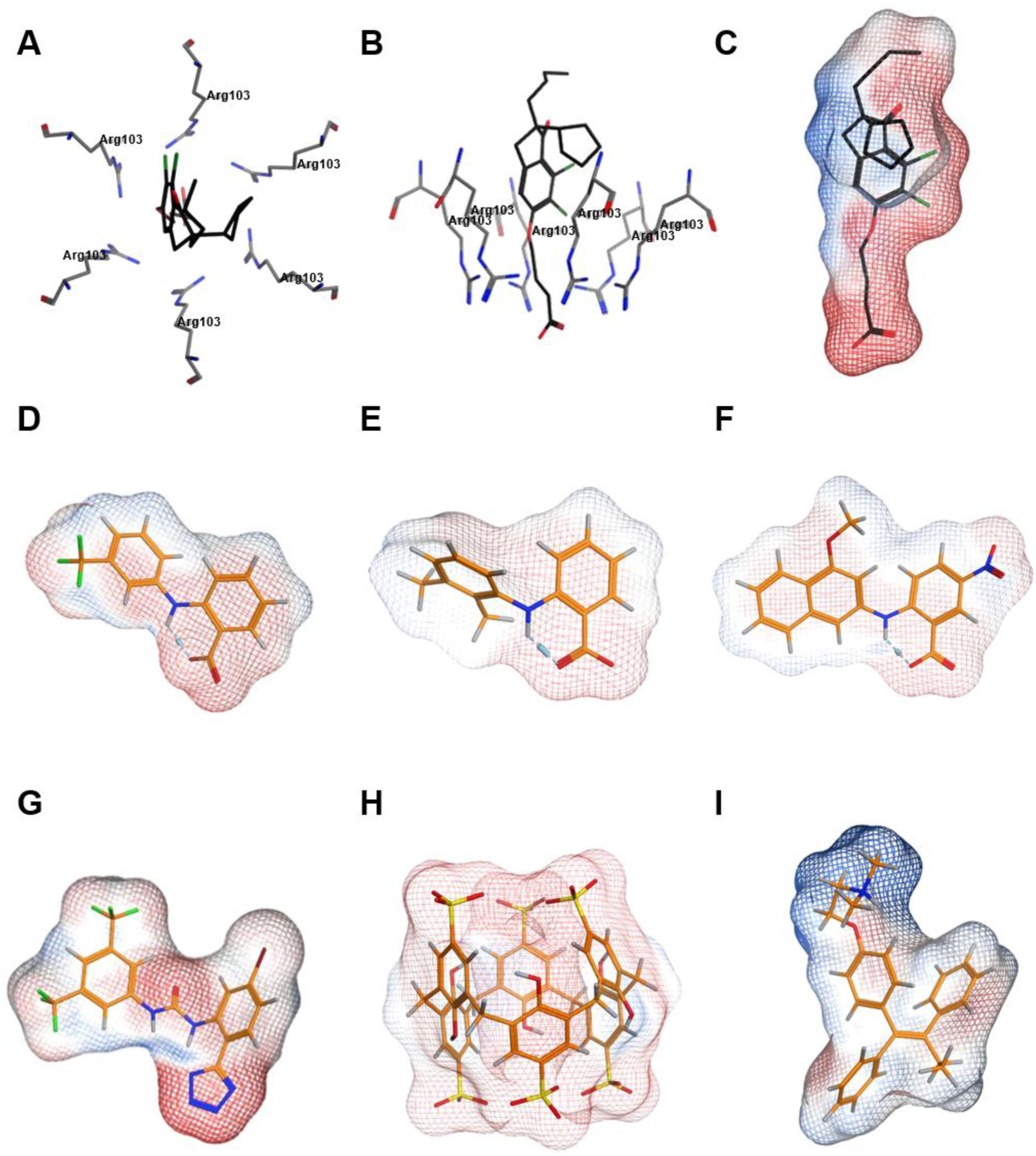
**(A)** Top view and **(B)** side view of the cryo-EM structure of DCPIB in the arginine pore of VRAC **(C-I)**. Molecular surfaces coloured by electrostatic potential for the docked poses of VRAC inhibitors, indicating regions of negative (red) and positive (blue) potential. **(C)** cryo-EM pose of DCPIB. Docked poses of **(D)** flufenamic acid **(E)** mefenamic acid **(F)** MONNA **(G)** NS3728 **(H)** 4-sulfonic calix(6)arene and **(I)** tamoxifen.

**Supplementary Figure 2.**
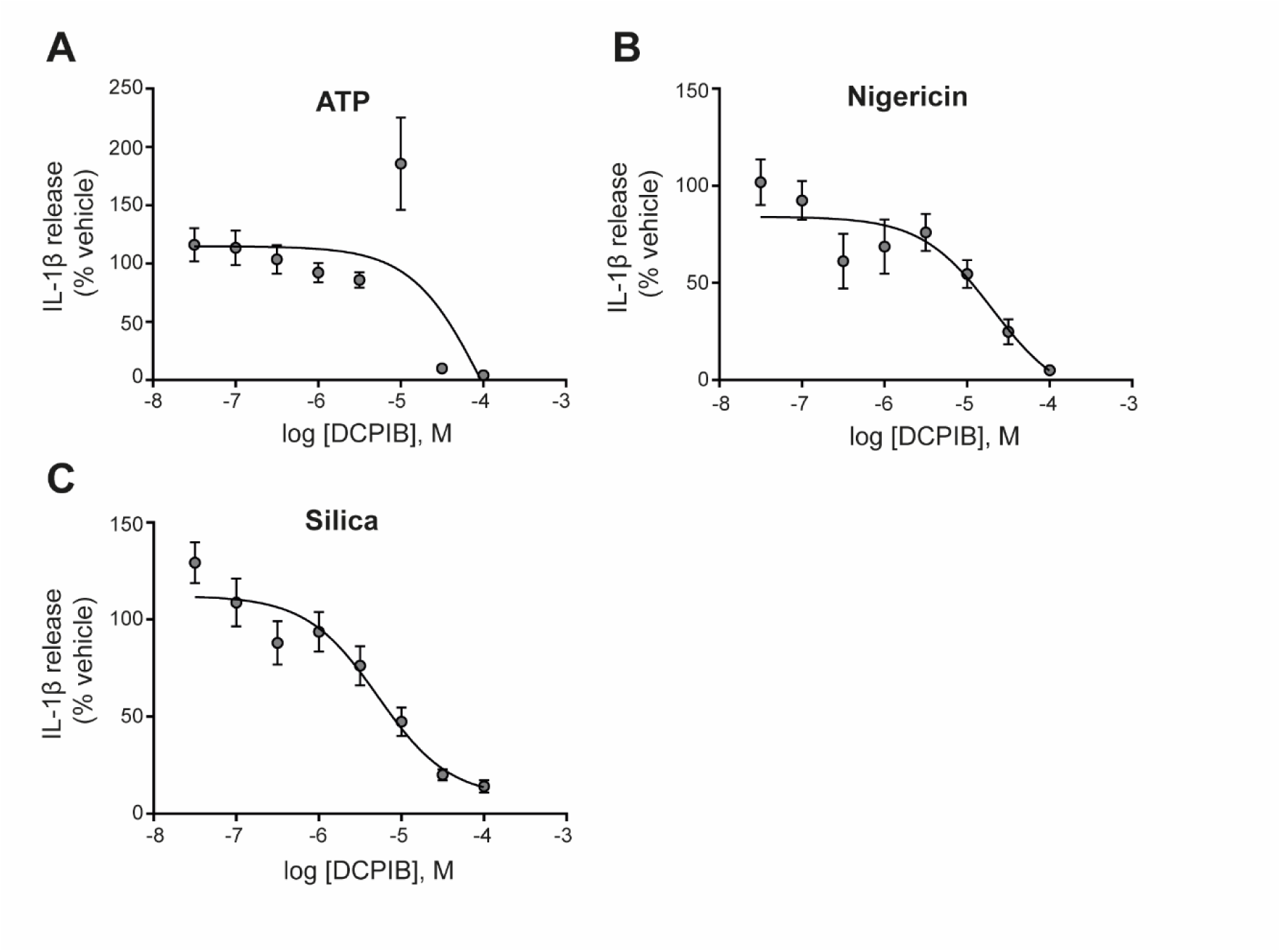
Dose-response curve for DCPIB. IL-1β release detected by ELISA on supernatants from LPS-primed (1 µg mL^-1^, 4 h) bone marrow derived macrophages (BMDM) pre-treated with the indicated dose of DCPIB (0.03-100 µM, 15 min) before stimulation with ATP (5 mM), nigericin (10 µM) or silica (300 µg mL^-1^) for 2 h (n=6). Experiments with different stimuli were performed in parallel. IL-1β release was normalised to that of vehicle (DMSO)-treated BMDMs. Dose-response curves were fitted using a three parameter logistical (3PL) model. Values shown are mean plus the SEM.

**Supplementary Figure 3.**
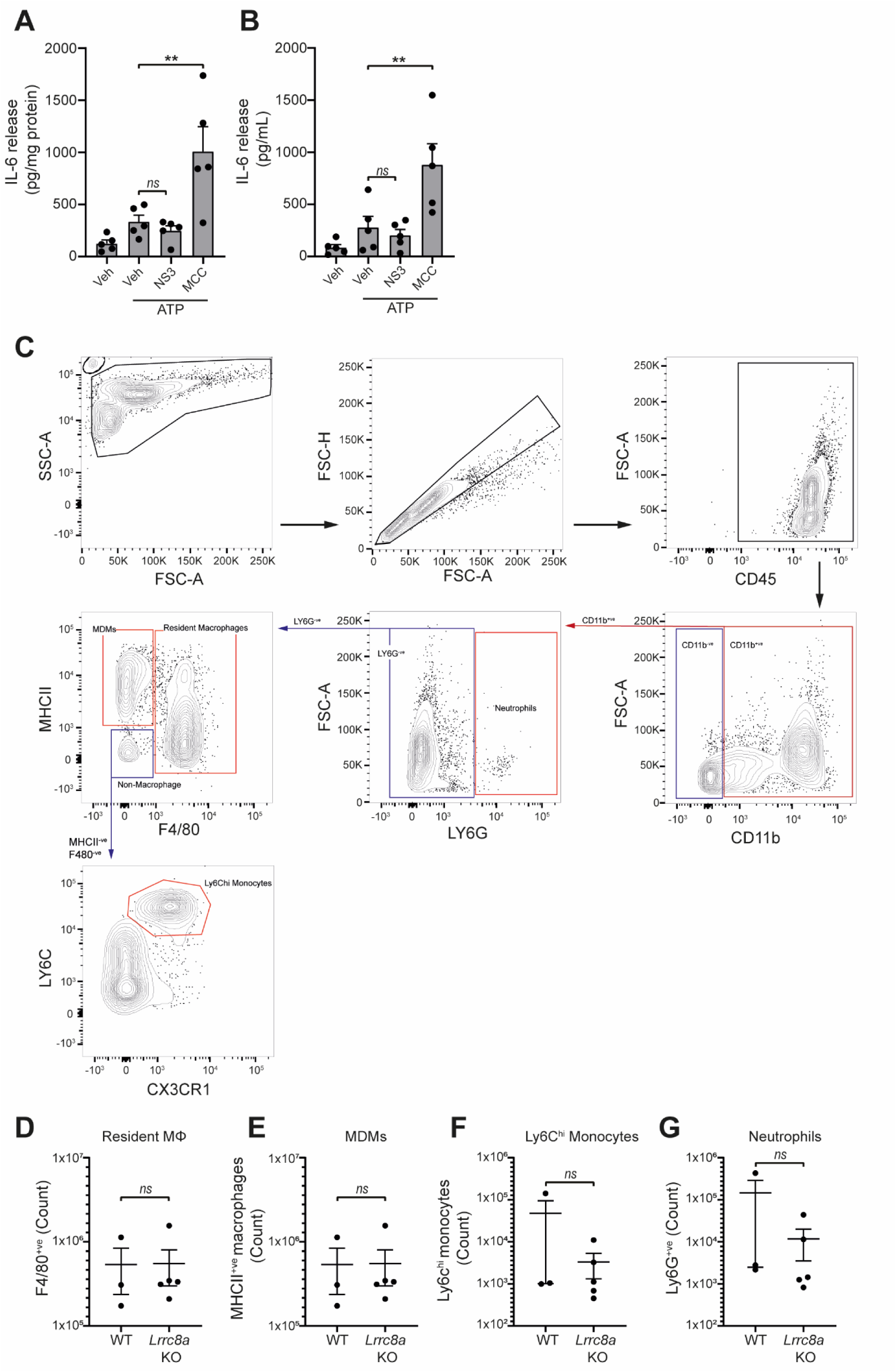
Loss of LRRC8a does not affect myeloid populations in the peritoneum. **(A-B)** IL-6 detected by ELISA in the peritoneal lavage **(A)** or plasma **(B)** from wild-type mice. Mice were pre-treated intraperitoneally (i.p.) with a vehicle control, NS3728 (NS3, 50 mg kg^-1^) or MCC950 (MCC, 50 mg kg^-1^) and LPS (1 µg). 4 h after injection with LPS, mice were anaesthetised and injected with additional vehicle control, NS3728 (NS3, 50 mg kg^-1^) or MCC950 (MCC, 50 mg kg^-1^) before i.p. injection of ATP (100 mM, 500 µL, 15 min) (n=5). NS3728 treatment did not significantly alter IL-6 levels. Injection of MCC950 significantly enhanced IL-6 release in the peritoneum and plasma compared to vehicle. **(C-G)** Flow cytometry of immune cells in naïve peritoneal lavage from WT or *Lrrc8a* KO mice (n=3-5). Representative gating strategy (C) and quantification of immune cells (D-G). Immune cells were initially gated on CD45^+ve^/CD11b^+ve^ cells and cell populations were identified as follows: neutrophils (LY6G^hi^), monocyte-derived-macrophages (MDMs) (LY6G^-ve^/MHCII^hi^/F4/80^-ve^), resident macrophages (Mϕ) (LY6G^-ve^/F4/80^hi^), and Ly6C^hi^ monocytes (LY6G^-ve^/MHCII^-ve^/F4/80^-ve^/CX3CR1^hi^/Ly6C^hi^). *ns* not significant, \*\**p*<0.01 determined by a one-way ANOVA with Dunnett’s (vs vehicle control) *post hoc* analysis (A,B) or an unpaired *t*-test (D-G). Values shown are mean plus the SEM.

**Supplementary video 1 – Phase contrast time-lapse of the regulatory volume decrease of wild-type (WT) littermate bone marrow-derived macrophages (BMDMs)**. WT BMDMs were incubated in an isotonic buffer (340 mOsm kg^-1^) for 5 minutes before dilution to a hypotonic solution (117 mOsm kg^-1^) for the duration of the recording. Images were captured every minute (n=3, Scale = 20 µm).

**Supplementary video 2 – Phase contrast time-lapse of the regulatory volume decrease of *Lrrc8a knockout* (KO) bone marrow-derived macrophages (BMDMs)**. *Lrrc8a KO* BMDMs were incubated in an isotonic buffer (340 mOsm kg^-1^) for 5 minutes before dilution to a hypotonic solution (117 mOsm kg^-1^) for the duration of the recording. Images were captured every minute (n=3, Scale = 20 µm).

